# Daclatasvir, a symmetric drug for an anti-symmetric target

**DOI:** 10.1101/2025.08.11.669771

**Authors:** Luis Córdova-Bahena, Axel Sánchez-Álvarez, Ivonne López-Lerma, Nohemí Salinas-Jazmín, José L. Medina-Franco, Marco Velasco-Velázquez

## Abstract

Immunotherapy has become a cornerstone in cancer treatment, with anti-PD-L1 antibodies effectively used across various cancers. Although these therapies have shown success, antibodies face limitations in bioavailability compared to low molecular mass compounds. An alternative strategy is to stabilize PD-L1 homodimers to prevent their immunosuppressive activity. The homodimer interface forms a tunnel-like cavity that can accommodate small molecules. However, no small drugs targeting PD-L1 homodimers have been approved for cancer treatment. Drug repurposing offers a promising approach to bridge this gap. In this study, we sought to identify potential PD-L1 inhibitors among FDA-approved drugs using virtual screening, followed by molecular docking, molecular dynamics simulations, and MM/PBSA binding energy calculations. Our results indicate that daclatasvir, an FDA-approved antiviral for hepatitis C, forms a stable and energetically favorable complex with the PD-L1 homodimer, suggesting it as a promising candidate for further investigation in cancer immunotherapy. Due to its symmetry, daclatasvir simultaneously interacts with both PD-L1 monomers in an equivalent manner, bridging the dimer interface. Its biphenyl core anchors at the center of the tunnel, the imidazole rings position at the entrances, and the pyrrolidine rings remain exposed to the solvent. Our in-depth characterization of the binding mode of daclatasvir clarifies its binding mechanism, and recent experimental findings have also indicated that daclatasvir binds to PD-L1, supporting its potential in this new context.

## 1. Introduction

Immune checkpoints are negative regulators of the immune system that maintain self-tolerance and control the intensity of immune responses [1]. The binding of Programmed Death-Ligand 1 (PD-L1) to Programmed Cell Death Protein 1 (PD-1) controls an immune checkpoint with key relevance in cancer [2]. PD-1 is expressed on T cells, whereas PD-L1 is present on both healthy and cancer cells. Under normal physiological conditions, PD-1 binds to PD-L1 triggers downstream signaling that attenuates T cell receptor and CD28 pathways, ultimately reducing T cell activation, proliferation, and cytokine production [3]. This deactivation is crucial to prevent excessive immune responses, thereby protecting tissues from autoimmune damage and maintaining immune tolerance. In the tumor microenvironment, cancer cells exploit this interaction to evade the immune response, allowing them to escape immune surveillance [4, 5]. PD-L1 has been reported as overexpressed in melanoma, thymoma, lung cancer, gastric cancer, hepatocellular carcinoma, renal cell carcinoma, esophageal cancer, pancreatic cancer, ovarian cancer, and bladder cancer [6–8]. Thus, PD-L1 has become an important clinically validated target in cancer therapy. Since 2016, three anti-PD-L1 antibodies have been approved for the treatment of various cancers [9]. Today, immunotherapy targeting PD-1/PD-L1 axis immune checkpoint is considered a successful approach in cancer care [10, 11].

Despite the success of anti-PD-L1 therapy, antibodies, as biotechnological drugs, are not exempt from adverse events and inherent toxicity [12]. Furthermore, they face challenges related to long half-life, immunogenicity, and limited permeability in tumor tissues [13]. Additionally, high costs limit the overall cost-effectiveness of these therapies in cancer treatment [14, 15]. Conversely, low molecular mass compounds offer notable advantages over monoclonal antibodies in addressing these issues. In recent years, various research groups have focused on developing small-molecule compounds with novel mechanisms that prevent PD-L1 from binding to its receptor by inducing PD-L1 homodimerization [16–18].

As a monomer, PD-L1 lacks a pocket to accommodate small compounds and is considered an undruggable target [19, 20]. However, when PD-L1 forms a homodimer, a hydrophobic pocket resembling a tunnel is created at the interface of the monomers. Further, crystallographic data deposited in the Protein Data Bank (PDB) [21] revealed a common biphenyl moiety from ligands inside this pocket, with polar substituents oriented toward the tunnel entrances and extending outward. Additionally, the PD-L1 homodimer exhibits an anti-symmetric arrangement with identical moieties on both sides of the tunnel. Because the PD-L1 homodimer is an anti-symmetric target, symmetry considerations are crucial in developing PD-L1 homodimer stabilizers [22]. On the other hand, C2 symmetry describes molecules with two-fold rotational symmetry, appearing identical after a 180° rotation around an axis. This structural insight has led to the development of several C2 symmetric biphenyl-based compounds [23]. For example, Basu *et al*., reported the design and evaluation of a symmetric ligand (**Figure 1**, ligand A) with PubChem CID: 138753643, which exhibited nanomolar activity in homogeneous time resolved fluorescence (HTRF) assays [24]. Another notable case is the ligand synthesized by Kawashita *et al*. (**Figure 1**, ligand B) with PubChem CID: 155536299, with micromolar activity in surface plasmon resonance (SPR) assay against PD-L1 [25]. Interestingly, both compounds are C2-symmetric stabilizers of PD-L1 homodimers. However, neither of these compounds has been approved as an antineoplastic drug, highlighting the need for additional compounds with similar activity.

**Figure. 1.**
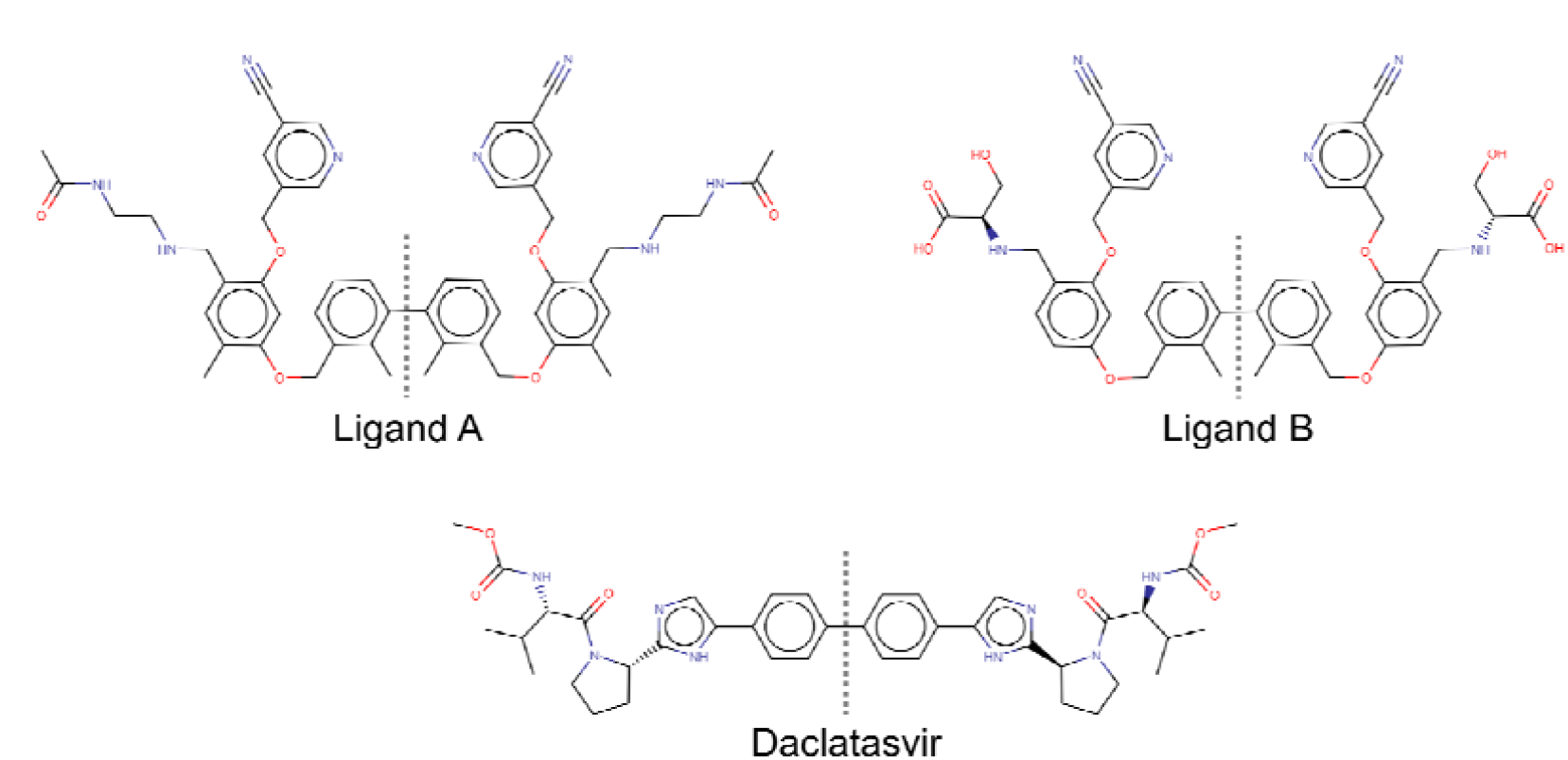
Chemical structures of the reference compounds and daclatasvir, with the C2 axis of symmetry indicated by a dotted line.

Drug repurposing has become a successful strategy for identifying new therapeutic uses for existing drugs beyond their original medical indications [26]. Historically, drug repurposing was largely a matter of serendipity [27]. However, it now involves systematic and rational approaches [28]. In particular, computational methods play a critical role in identifying potential candidates for repurposing [29–31]. For instance, *in silico* high-throughput molecular docking and molecular dynamics (MD) simulations are computational techniques that provide valuable information to guide experimental studies in drug repurposing campaigns [32, 33].

Daclatasvir (**Figure 1**) is an antiviral drug used to treat hepatitis C virus infections and has been approved by the Food and Drug Administration (FDA) of the United States since 2015 [34]. From a structural perspective, daclatasvir is a C2-symmetrical compound with a biphenyl core and polar substituents on both sides. Summarizing, all three chemical structures —those of the two selected PD-L1 homodimer stabilizers and daclatasvir— exhibit C2-symmetry, featuring a central biphenyl moiety with polar substituents at both ends.

In this study, we performed high-throughput molecular docking of the DrugBank database into the PD-L1 homodimer, aiming to identify potential stabilizers. We hypothesized that FDA-approved drugs with C2 symmetry and a hydrophobic central core could act as PD-L1 stabilizers. Daclatasvir achieved a top-20 docking score (**Table S1**) and was the only compound in this group with C2 symmetry. Daclatasvir has the structural features for forming stacking interactions with PD-L1 tyrosine residues and substituents with multiple hydrogen bond acceptors and donors, creating a central hydrophobic core flanked by polar regions that complement the features of the PD-L1 tunnel. Then, we characterized the complexes of the PD-L1 homodimer with daclatasvir, or each of two reference compounds [24, 25] through molecular dynamics (MD) simulations. Binding energies for all three compounds were calculated using the molecular mechanics Poisson–Boltzmann surface area (MM-PBSA) approach [35]. Our findings show that daclatasvir forms a stable and energetically favorable complex with the PD-L1 homodimer, suggesting it is a promising candidate for repurposing as a PD-L1 homodimer stabilizer. Notably, during the course of this investigation, it was published that daclatasvir indeed is able to bind human PD-L1 [36].

## 2. Methodology

### 2.1 Molecular docking

High-throughput molecular docking was conducted using Molecular Operating Environment (MOE) software v2023.10. The receptor structure, co-crystallized with ligand A, was obtained from the PDB with ID 6RPG [24]. One monomer was tagged as PD-L1_A_ and the other one as PD-L1_B_. Ligand B was obtained from the PubChem database [37]. Compounds from the DrugBank database [38], along with reference ligands A and B, were imported into MOE. All ligands were protonated at pH 6.7 to simulate the extratumoral environment [39] and optimized using the MMFF94x force field. All water molecules were removed. Initial ligand poses were generated using the Proxy Triangle placement method and evaluated with the London dG scoring function. Docking accuracy was validated by redocking the co-crystallized ligand A in the structure, following the protocol described above.

### 2.2 Molecular dynamics simulation

Structures of the complexes were obtained through molecular docking as described above. The systems were prepared using CHARMM-GUI [40], where the protein was processed for MD simulation. Preparation steps included the addition of missing hydrogen atoms, assignment of protonation states at pH 6.4-7.1, and parameterization using the CHARMM36m force field [41]. Both nitrogen atoms of the imidazole side chain in all histidine residues were protonated. The protein was solvated in a cubic box of 85 Å by side with TIP3P water molecules, with counterions added to neutralize the system and achieve a physiological ionic strength of 0.15 M. The prepared system was imported into GROMACS v2021.6 [42] and subjected to energy minimization using the steepest descent algorithm. The system then underwent equilibration in two phases: first, in the NVT ensemble, maintaining the temperature at 310.15 K using the modified Berendsen thermostat; and second, in the NPT ensemble, with the pressure maintained at 1 atm using the Parrinello-Rahman barostat. Finally, a production MD simulation was performed for 200 ns with a time step of 2 fs. Periodic boundary conditions were applied, and the LINCS algorithm was used to constrain bond lengths involving hydrogen atoms. Coordinates were saved every 10 ps for subsequent analysis.

### 2.3 Binding affinity

The binding affinity of the reference and daclatasvir complexes was assessed using MM/PBSA approach, based on 30 energetically accessible conformations extracted from the last 30 ns of MD trajectories [43].

## 3. Results

Building on recent findings that daclatasvir, a C2-symmetric molecule with a biphenyl core, binds to the PD-L1 homodimer, this study seeks to shed light on its detailed binding interactions and mechanisms of stabilization.

### 3.1 Daclatasvir is predicted to bind the PD-L1 homodimer

First, we redocked the co-crystallized ligand A into the PD-L1 homodimer structure. The calculated pose was compared with the crystallographic conformation, yielding a root mean square deviation (RMSD) of 1.90 Å (**Figure S1**). Second, the binding modes of the symmetric ligand B reported by Kawashita *et al*. and daclatasvir were identified using the validated docking protocol focusing on the top-ranked scored poses.

Both ligands A, and B belong to the class of biphenyl-based compounds, although they contain a more complex tetraphenyl core. Ligand A has two identical substituents on each side of the tetraphenyl core. These are a cyanopyridine and a charged polar chain. An analysis of the PD-L1 homodimer/ligand A complex showed that these substituents interacted with both PD-L1 monomers. The cyanopyridine and positively charged polar chain on one side bind to the PD-L1_A_ monomer, while the other set interacted with the PD-L1_B_ monomer (**Figure 2 A**).

**Figure 2.**
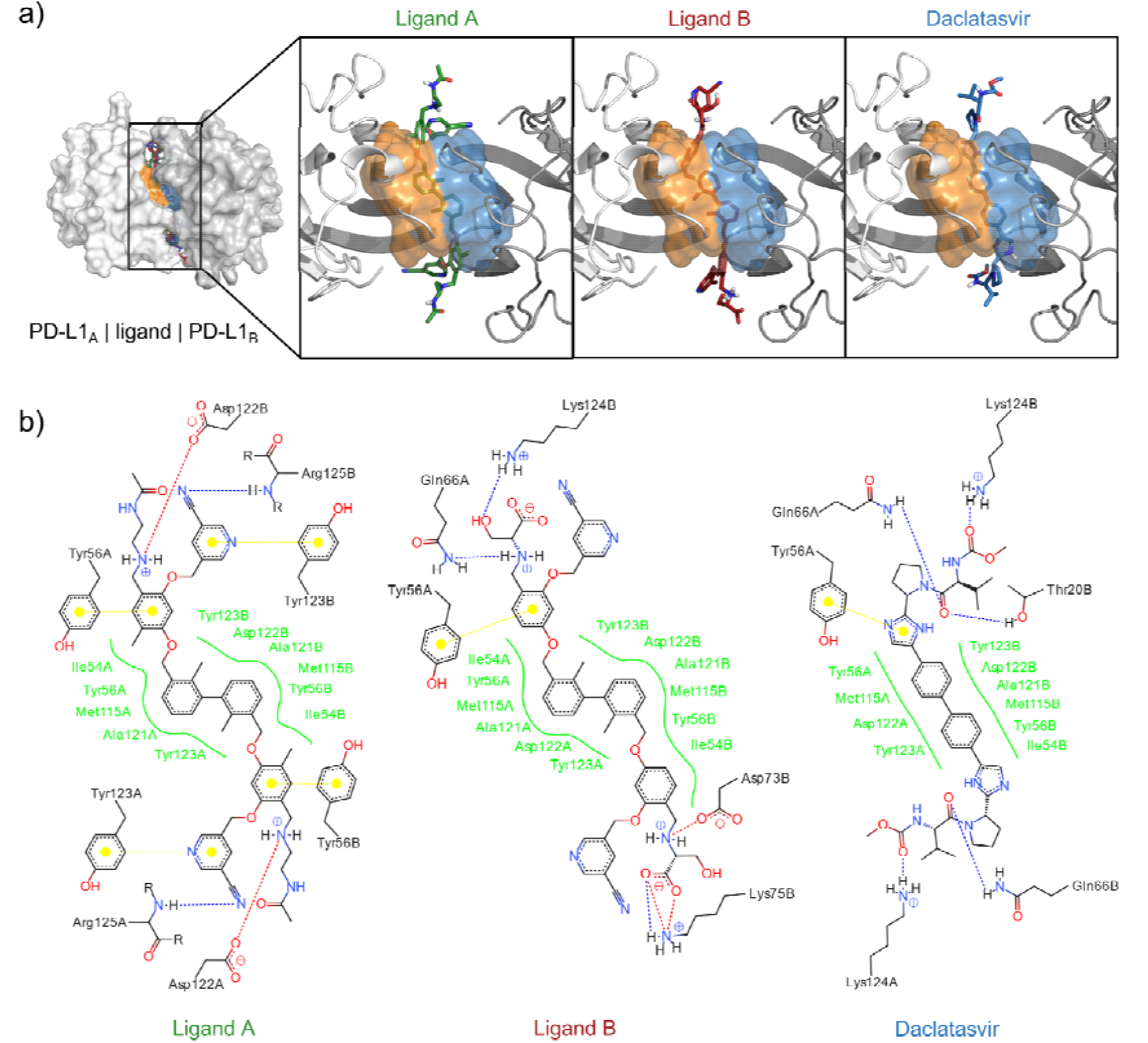
Binding modes for reference ligands plus daclatasvir. **a)** 3D structure of the complexes, showing two PD-L1 monomers, in surface representation, and the ligands in licorice. The insets highlight the biphenyl core and the orientations of the polar chains for each ligand. **b)** Intermolecular interactions for each ligand. Residues involved in hydrophobic interactions are shown in green. Stacking interactions, salt bridges, and hydrogen bonds are represented by dotted lines in yellow, red, and blue, respectively.

As expected, ligand B binding to the PD-L1 homodimer was mediated by its biphenyl core interacting with the center of the tunnel formed at the interface of the PD-L1 monomers. The two substituents on each side of the tetraphenyl core of ligand B are a cyanopyridine and a double-charged polar chain. The cyanopyridine in ligand B exhibited no intermolecular interactions with either monomer. However, one of the two double-charged chains was oriented toward the PD-L1_B_ monomer while the other interacts with both monomers (**Figure 2 A**).

As was hypothesized, daclatasvir was bound to the PD-L1 homodimer with the biphenyl core positioned in the center of the tunnel. In addition, daclatasvir has two aromatic rings as substituents on each side of the biphenyl core. These aromatic rings were located at the entrances of the tunnel.

Moreover, the polar moieties attached to the extra rings extend outside the tunnel and remain away from the core (**Figure 2A**).

An in-depth analysis of intermolecular interactions was performed to identify key residues involved in ligand binding. For all three ligands, the binding mode was primarily driven by hydrophobic interactions with residues located in the center of the tunnel, such as Ile54, Tyr56, Met115, Ala121, and Tyr123 from both monomers. Additionally, stacking interactions with Tyr 56 were observed at the tunnel entrance. For ligand A, stacking occurred with both monomers, while for ligand B and daclatasvir, it appeared only with one monomer. Ligand A also formed extra stacking interactions with Tyr123 from both monomers. Furthermore, salt bridges were formed between the positively charged moieties of ligand A and ligand B and negatively charged residues Asp122 (both monomers) and Asp75B, respectively. For ligand B, salt bridges were observed between its negatively charged moiety and the positively charged residues Lys124 from both monomers, while daclatasvir formed hydrogen bonds with these residues. Additionally, daclatasvir established additional hydrogen bonds with Thr20B and Gln66 from both monomers. Ligand A formed hydrogen bonds with the main chain of Arg 125 from both monomers, while ligand B interacted with the side chains of Gln66A and Lys124B (**Figure 2B**). These findings indicate a common binding pattern among ligand A, ligand B, and daclatasvir.

### 3.2 Daclatasvir forms a stable complex with the PD-L1 homodimer

Molecular dynamics (MD) studies were performed to evaluate the stability of the daclatasvir/PD-L1 complexes. Complexes with ligands A, and B were used as positive controls. As expected, the PD-L1 homodimer/ligand A complex showed high stability, with an RMSD of 2.31 ± 0.26 Å for the ligand and 1.84 ± 0.20 Å for the protein backbone (**Figure 3A**). Non-relevant conformational changes from the pose obtained previously by crystallography were observed for the Ligand A (**Figure 3D**). On the other hand, the RMSD for ligand B (5.86 ± 0.59 Å) indicated this ligand adopted a new conformation (**Figure. 3B**). The shift was primarily caused by the reorientation of the biphenyl core substituents into a position opposite to that predicted by molecular docking (**Figure 3E**).

**Figure 3.**
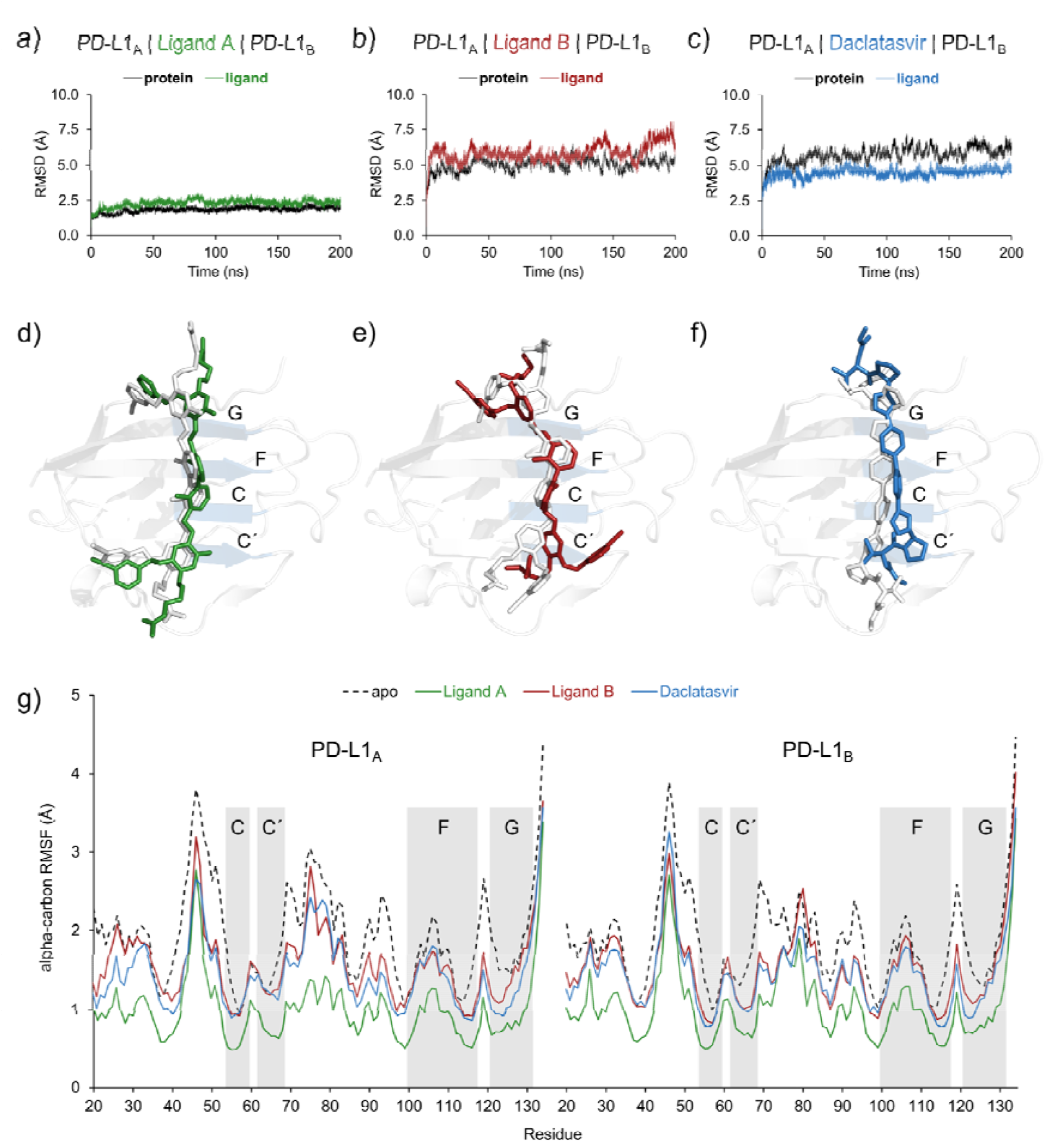
Dynamic behavior of reference ligands and daclatasvir, demonstrating their stabilizing effects on the PD-L1 homodimer. **a-c)** RMSD plots of the PD-L1 homodimer bound to **a)** ligand A, **b)** ligand B, or **c)** daclatasvir. **d-f)** Representative bioactive conformations obtained from MD simulations for **d)** ligand A (green), **e)** ligand B (red), and **f)** daclatasvir (blue). The initial conformation of ligands is shown in white-colored licorice model. Only the structure of the PD-L1_B_ monomer, represented in a cartoon model, is shown for clarity. Relevant beta sheets are labeled. **g)** RMSF of the apo PD-L1 homodimer (dashed line) compared to the complexes with ligand A, ligand B, and daclatasvir (colored in green, red, and blue, respectively). Beta sheets are shown in gray and labeled accordingly.

For the PD-L1 homodimer/Daclatasvir complex, an RMSD of 4.50 ± 0.33 Å was observed for the ligand, indicating that daclatasvir also adopts a new conformation (**Figure 3C**). In contrast to ligands A, and B, the biphenyl core of daclatasvir shifted into a significantly different position (**Figure 3F**). However, this new conformation remained stable. The RMSD values for the PD-L1 homodimer in complexes with ligand B or daclatasvir were 5.06 ± 0.45 Å, and 5.75 ± 0.54 Å, respectively indicating no significant conformational changes in the protein (**Figure 3B, 3C**).

To assess the effect of the ligands on PD-L1 homodimer, root mean square fluctuation (RMSF) values were calculated for the three complexes described above. The presence of the ligands reduced residue fluctuation compared to the apo PD-L1 homodimer. The complex with ligand A exhibited the lowest overall fluctuation. In particular, the C and C’ beta sheets showed minimal movement, though the interconnection loop between them remained flexible. A similar trend was observed for the terminal region of the F beta sheet and the beginning of the G beta sheet. As expected for ligand B and hypothesized for daclatasvir, these systems showed a similar pattern of reduced fluctuation compared to the apo homodimer. Notably, the interconnection loop of the C and C’ beta sheets in the daclatasvir complex exhibited less fluctuation compared to ligand B (**Figure 3G**). These findings indicate that ligand A, ligand B, and daclatasvir stabilize the PD-L1 homodimer in a similar way.

### 3.3 Daclatasvir shows comparable binding affinity to reported PD-L1 inhibitors

To evaluate the affinity to PD-L1 homodimer, the binding energies of ligands A, B, and daclatasvir were calculated from three replicates each, using the MM/PBSA approach (**Table S2, Figure 4a**). As expected for ligands A and B and hypothesized for daclatasvir, the binding energy was favorable. We found no statistically significant differences between ligands in the calculated ΔGs. However, ΔH was significantly more favorable for ligands A and B than for daclatasvir, with ligand A having almost double the ΔH of daclatasvir. Despite this, the ΔS contribution was better for daclatasvir binding. The entropy change for ligands A and B was three times higher than for daclatasvir, likely due to the greater number of accessible bioactive conformations. Ligands A and B each have 23 rotatable bonds, while daclatasvir has only 13, limiting its conformational flexibility.

**Figure 4.**
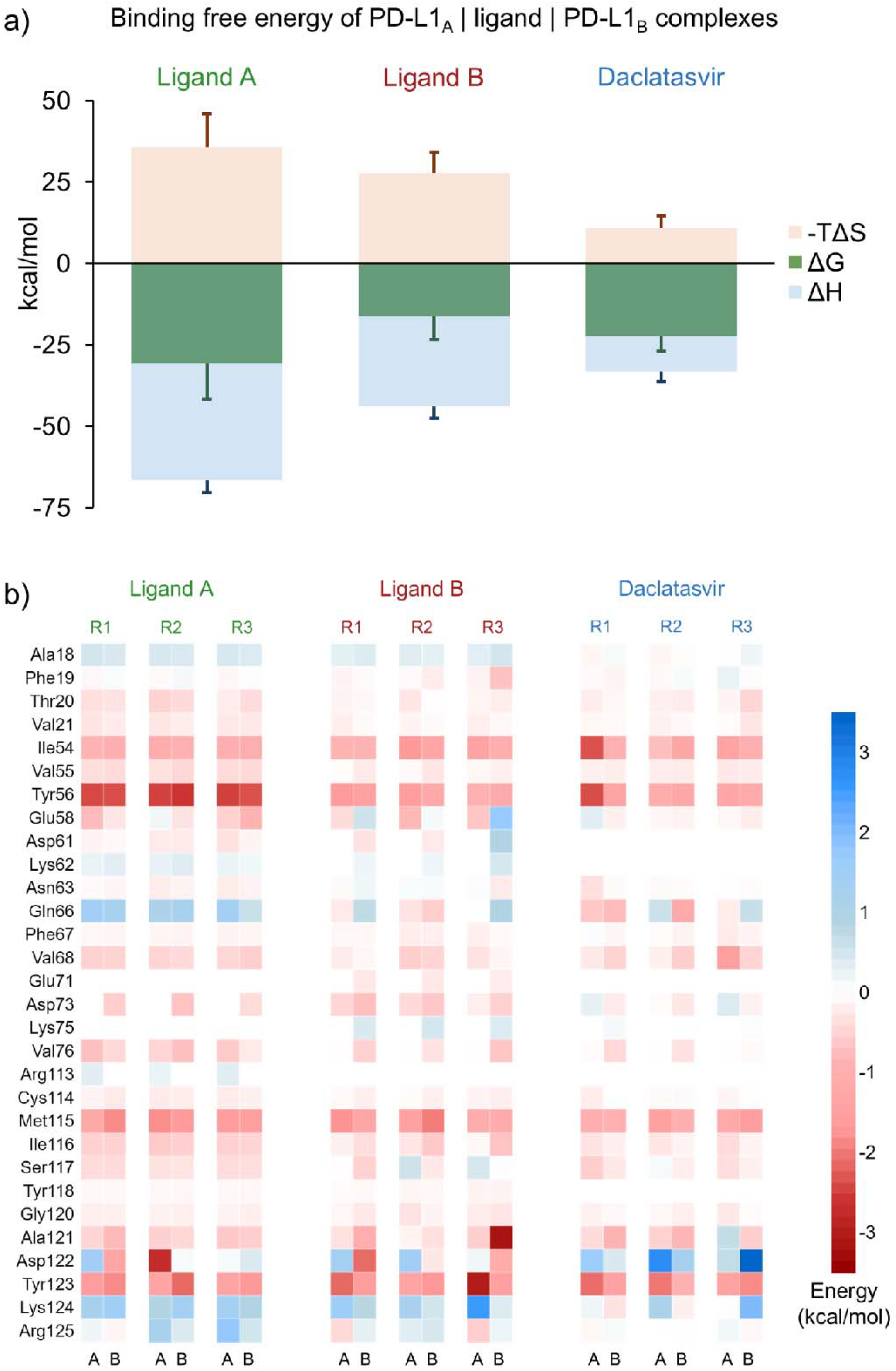
Total binding energy, and contribution by residue. **a)** Change in Gibbs free energy (ΔG) with enthalpy (ΔH) and entropy (ΔS) changes contributions, and **b)** Per-residue contribution to binding energy for complexes PD-L1 homodimer-ligand A,-B, and-daclatasvir. T means temperature.

To explore in detail the binding modes of ligands A, B, and daclatasvir, we performed a residue-wise binding energy analysis (**Figure 4b**). The tunnel residues, Tyr56 at the entrance and exit, and Met115 in the middle, were key contributors for all ligands. Additionally, Ile54, Val55, Gly120, and Ala121 at the extremes of the tunnel contributed favorably in a similar manner for all three ligands. Tyr123 from both monomers also played a central role in binding for all three ligands. On the other hand, residues outside of the targeted tunnel showed minimal contributions for all ligands. For example, Ala18 from both monomers contributed unfavorably to ligand A and B binding. However, for daclatasvir, Ala18 from monomer A contributed favorably, while Ala 18 from monomer B had an unfavorable impact. Phe19 from each monomer contributed differently in ligands A and daclatasvir, but contributed favorably in ligand B. Other residues in the N-terminal loop, such as Thr20, Val21, and Thr22, had small yet favorable and symmetric contributions for all ligands.

We also identified other residues that mediate ligand-or monomer-specific interactions. Asp122 had a mixed role; for ligands A and B it contributed favorably from one monomer and unfavorably from the other. On the other hand, it showed unfavorable energy for both monomers in the case of daclatasvir. Gln66 contributed unfavorably for ligand A and favorably for daclatasvir, while for ligand B, it had opposite roles depending on the monomer analyzed. All results were consistent through three replicates. This exhaustive characterization of the binding modes of reference ligands A, and B shows a highly symmetrical contribution of binding energy from each PD-L1 monomer. Finally, and most importantly, our findings show that daclatasvir emulates the binding mode of reported inhibitors with comparable affinity, primarily due to reduced entropic effects.

## 4. Discussion

Daclatasvir, an antiviral drug approved for HCV, acts by inhibiting RNA replication and virion assembly through binding to NS5A, a nonstructural HCV phosphoprotein. Since the crystal structure of NS5A remains unresolved, consensus on the exact binding site for daclatasvir is lacking. However, an NS5A homology model was developed, followed by molecular docking and MD studies to elucidate the daclatasvir binding mode. This model suggests that daclatasvir binds symmetrically to the NS5A homodimer, which constitutes the biological unit [44]. These findings suggest the potential of daclatasvir to bind to anti-symmetrical homodimers and highlights the importance of the symmetry on its structure.

Herein, we have provided a solid basis by computational modeling that daclatasvir has the potential to bind to the PD-L1 homodimer. The MD simulations and binding energy calculations suggest that daclatasvir forms a stable complex with the PD-L1 homodimer. We hypothesize that other symmetric molecules, including symmetric stereoisomers and *meso* forms, can target both the PD-L1 and NS5A homodimers. Increasing the rigidity of molecules by reducing the rotatable bonds may improve the entropic contribution. Recent studies by Sun *et al*. have reported the experimental binding of daclatasvir to human PD-L1 in HepG2 and Jurkat cells [36]. Additionally, daclatasvir has been shown to increase T-cell levels in patients undergoing antiviral treatment for chronic hepatitis C [45]. It has been reported that PD-L1 homodimerization promotes its internalization and degradation [46, 47]. Furthermore, direct-acting antiviral treatments, including daclatasvir, have been reported to downregulate immune checkpoint expression [48].

Taken together, these findings suggest that daclatasvir holds potential as a candidate for immune checkpoint inhibition. Stabilizing PD-L1 homodimers could prevent cancer cells from evading immune surveillance, which is particularly relevant given the increasing interest in small-molecule alternatives to monoclonal antibodies for targeting immune checkpoints. For example, BMS-103 and BMS-142 bind strongly to the PD-L1 homodimer, preventing the formation of the PD-1/PD-L1 axis and promoting T cell function. However, their immunological efficacy is compromised due to their acute cytotoxicity [49]. Another notable example is the compound Incyte-011, which binds to the PD-L1 homodimer and increases IFN-γ production. However, it also exhibits high cytotoxicity [50]. In contrast, as an approved drug, daclatasvir offers significant advantages in terms of clinical development and safety profiling.

Our MM/PBSA computations indicate the following affinity ranking: ligand A> ligand B> daclatasvir. These results cannot be directly compared with the available experimental data, since the assays for evaluation of ligand binding were different. Ligand A showed an IC□□ of 3.0 nM in an HTRF assay, while ligand B and daclatasvir exhibited K_D_ values of 0.019 nM and 11.4 μM, respectively, in a SPR assays [24,25,36]. These methodological differences and reported parameters highlight the challenge of directly correlating their affinities. Additionally, it is known that the predictions made based on computational analysis require further experimental validation [51–53]. Thus, it is necessary to assess daclatasvir’s efficacy and specificity in relevant models of disease including animal models. Such experimental evaluations could confirm the activity of daclatasvir as a PD-L1 homodimer stabilizer and would support its repurposing as an antineoplastic agent.

## 5. Conclusions

Overall, our study provides a compelling case for the potential of daclatasvir as a PD-L1 homodimer stabilizer. High-throughput molecular docking identified daclatasvir, a C2-symmetric compound with a biphenyl core, as a top-ranking candidate. Furthermore, Sun *et al*. reported that daclatasvir binds to PD-L1, and our molecular dynamics simulations and binding energy calculations offered deeper insights into this interaction. Interestingly, MM/PBSA analysis revealed that daclatasvir demonstrated a minimal, unfavorable ΔS contribution compared to reference ligands, highlighting the key role of entropy in binding affinity. These findings underscore the need for further exploration of the mechanism by which daclatasvir acts and its potential therapeutic applications in cancer immunotherapy.

## Supporting information

Supplementary material of Daclatasvir, a symmetric drug for an anti-symmetric target

## 6. Acknowledgements

This research was supported by CONAHCYT 319300 and LANDCAD-UNAM-DGTIC-386 (L. C-B.); Cátedras CONAHCYT 639 (L. C-B. and M. V-V.); and CONAHCYT fellowship 1309789 (A. S-A.).

## 7. Authors’ contributions

L.C-B. and M. V-V. conceived and designed the study. A. S-A., and I. L-L. performed the computational experiments. L. C-B. and A. S-A. analyzed the data and contributed to the interpretation of the results. L. C-B. was responsible for writing the manuscript, with critical revisions and intellectual input from M. V-V., N. S-J, and J. M-F.. All authors read and approved the final version of the manuscript.

## References

1. Zhang, Y., Zheng, J.: Functions of Immune Checkpoint Molecules Beyond Immune Evasion. In: Xu, J. (ed.) Regulation of Cancer Immune Checkpoints: Molecular and Cellular Mechanisms and Therapy. pp. 201–226. Springer, Singapore (2020)

2. Zak, K.M., Grudnik, P., Magiera, K., Dömling, A., Dubin, G., Holak, T.A.: Structural Biology of the Immune Checkpoint Receptor PD-1 and Its Ligands PD-L1/PD-L2. Structure. 25, 1163–1174 (2017). 10.1016/j.str.2017.06.011

3. Hui, E., Cheung, J., Zhu, J., Su, X., Taylor, M.J., Wallweber, H.A., Sasmal, D.K., Huang, J., Kim, J.M., Mellman, I., Vale, R.D.: T cell costimulatory receptor CD28 is a primary target for PD-1–mediated inhibition. Science. 355, 1428–1433 (2017). 10.1126/science.aaf1292

4. Marcucci, F., Rumio, C., Corti, A.: Tumor cell-associated immune checkpoint molecules – Drivers of malignancy and stemness. Biochimica et Biophysica Acta (BBA) - Reviews on Cancer. 1868, 571–583 (2017). 10.1016/j.bbcan.2017.10.006

5. Jiang, X., Wang, J., Deng, X., Xiong, F., Ge, J., Xiang, B., Wu, X., Ma, J., Zhou, M., Li, X., Li, Y., Li, G., Xiong, W., Guo, C., Zeng, Z.: Role of the tumor microenvironment in PD-L1/PD-1-mediated tumor immune escape. Mol Cancer. 18, 10 (2019). 10.1186/s12943-018-0928-4

6. Patel, S.P., Kurzrock, R.: PD-L1 Expression as a Predictive Biomarker in Cancer Immunotherapy. Molecular Cancer Therapeutics. 14, 847–856 (2015). 10.1158/1535-7163.MCT-14-0983

7. Ullah, A., Pulliam, S., Karki, N.R., Khan, J., Jogezai, S., Sultan, S., Muhammad, L., Khan, M., Jamil, N., Waheed, A., Belakhlef, S., Ghleilib, I., Vail, E., Heneidi, S., Karim, N.A.: PD-L1 Over-Expression Varies in Different Subtypes of Lung Cancer: Will This Affect Future Therapies? Clinics and Practice. 12, 653–671 (2022). 10.3390/clinpract12050068

8. Wang, X., Teng, F., Kong, L., Yu, J.: PD-L1 expression in human cancers and its association with clinical outcomes. OncoTargets and Therapy. 9, 5023–5039 (2016). 10.2147/OTT.S105862

9. Jiang, M., Liu, M., Liu, G., Ma, J., Zhang, L., Wang, S.: Advances in the structural characterization of complexes of therapeutic antibodies with PD-1 or PD-L1. mAbs. 15, 2236740 (2023). 10.1080/19420862.2023.2236740

10. Esfahani, K., Roudaia, L., Buhlaiga, N., Del Rincon, S.V., Papneja, N., Miller, W.H.: A review of cancer immunotherapy: from the past, to the present, to the future. Curr Oncol. 27, S87–S97 (2020). 10.3747/co.27.5223

11. Farkona, S., Diamandis, E.P., Blasutig, I.M.: Cancer immunotherapy: the beginning of the end of cancer? BMC Medicine. 14, 73 (2016). 10.1186/s12916-016-0623-5

12. Naidoo, J., Page, D.B., Li, B.T., Connell, L.C., Schindler, K., Lacouture, M.E., Postow, M.A., Wolchok, J.D.: Toxicities of the anti-PD-1 and anti-PD-L1 immune checkpoint antibodies. Annals of Oncology. 26, 2375–2391 (2015). 10.1093/annonc/mdv383

13. Zhan, M.-M., Hu, X.-Q., Liu, X.-X., Ruan, B.-F., Xu, J., Liao, C.: From monoclonal antibodies to small molecules: the development of inhibitors targeting the PD-1/PD-L1 pathway. Drug Discovery Today. 21, 1027–1036 (2016). 10.1016/j.drudis.2016.04.011

14. Liu, G., Kang, S., Wang, X., Shang, F.: Cost-Effectiveness Analysis of Atezolizumab Versus Chemotherapy as First-Line Treatment for Metastatic Non-Small-Cell Lung Cancer With Different PD-L1 Expression Status. Front. Oncol. 11, (2021). 10.3389/fonc.2021.669195

15. Hu, X., Hay, J.W.: First-line pembrolizumab in PD-L1 positive non-small-cell lung cancer: A cost-effectiveness analysis from the UK health care perspective. Lung Cancer. 123, 166–171 (2018). 10.1016/j.lungcan.2018.07.012

16. Bailly, C., Vergoten, G.: Protein homodimer sequestration with small molecules: Focus on PD-L1. Biochemical Pharmacology. 174, 113821 (2020). 10.1016/j.bcp.2020.113821

17. Liu, L., Zhang, H., Hou, J., Zhang, Y., Wang, L., Wang, S., Yao, Z., Xie, T., Wen, X., Xu, Q., Dai, L., Feng, Z., Zhang, P., Wu, Y., Sun, H., Liu, J., Yuan, H.: Discovery of Novel PD-L1 Small-Molecular Inhibitors with Potent In Vivo Anti-tumor Immune Activity. J. Med. Chem. 67, 4977–4997 (2024). 10.1021/acs.jmedchem.4c00102

18. Cheng, B., Wang, W., Liu, T., Cao, H., Pan, W., Xiao, Y., Liu, S., Chen, J.: Bifunctional small molecules targeting PD-L1/CXCL12 as dual immunotherapy for cancer treatment. Sig Transduct Target Ther. 8, 1–3 (2023). 10.1038/s41392-022-01292-5

19. Gashaw, I., Ellinghaus, P., Sommer, A., Asadullah, K.: What makes a good drug target? Drug Discovery Today. 16, 1037–1043 (2011). 10.1016/j.drudis.2011.09.007

20. Makley, L.N., Gestwicki, J.E.: Expanding the Number of ‘Druggable’ Targets: Non-Enzymes and Protein–Protein Interactions. Chemical Biology & Drug Design. 81, 22–32 (2013). 10.1111/cbdd.12066

21. Berman, H.M., Westbrook, J., Feng, Z., Gilliland, G., Bhat, T.N., Weissig, H., Shindyalov, I.N., Bourne, P.E.: The Protein Data Bank. Nucleic Acids Research. 28, 235–242 (2000). 10.1093/nar/28.1.235

22. Klimek, J., Kruc, O., Ceklarz, J., Kamińska, B., Musielak, B., van der Straat, R., DLJmling, A., Holak, T.A., Muszak, D., Kalinowska-Tłuścik, J., Skalniak, Ł., Surmiak, E.: C2-Symmetrical Terphenyl Derivatives as Small Molecule Inhibitors of Programmed Cell Death 1/Programmed Death Ligand 1 Protein–Protein Interaction. Molecules. 29, 2646 (2024). 10.3390/molecules29112646

23. Chen, R., Yuan, D., Ma, J.: Advances of Biphenyl Small-Molecule Inhibitors Targeting PD-1/PD-L1 Interaction in Cancer Immunotherapy. Future Medicinal Chemistry. 14, 97–113 (2022). 10.4155/fmc-2021-0256

24. Basu, S., Yang, J., Xu, B., Magiera-Mularz, K., Skalniak, L., Musielak, B., Kholodovych, V., Holak, T.A., Hu, L.: Design, Synthesis, Evaluation, and Structural Studies of C2-Symmetric Small Molecule Inhibitors of Programmed Cell Death-1/Programmed Death-Ligand 1 Protein–Protein Interaction. J. Med. Chem. 62, 7250–7263 (2019). 10.1021/acs.jmedchem.9b00795

25. Kawashita, S., Aoyagi, K., Yamanaka, H., Hantani, R., Naruoka, S., Tanimoto, A., Hori, Y., Toyonaga, Y., Fukushima, K., Miyazaki, S., Hantani, Y.: Symmetry-based ligand design and evaluation of small molecule inhibitors of programmed cell death-1/programmed death-ligand 1 interaction. Bioorganic & Medicinal Chemistry Letters. 29, 2464–2467 (2019). 10.1016/j.bmcl.2019.07.027

26. Pushpakom, S., Iorio, F., Eyers, P.A., Escott, K.J., Hopper, S., Wells, A., Doig, A., Guilliams, T., Latimer, J., McNamee, C., Norris, A., Sanseau, P., Cavalla, D., Pirmohamed, M.: Drug repurposing: progress, challenges and recommendations. Nat Rev Drug Discov. 18, 41–58 (2019). 10.1038/nrd.2018.168

27. Jourdan, J.-P., Bureau, R., Rochais, C., Dallemagne, P.: Drug repositioning: a brief overview. Journal of Pharmacy and Pharmacology. 72, 1145–1151 (2020). 10.1111/jphp.13273

28. Bellera, C.L., Di Ianni, M.E., Sbaraglini, M.L., Castro, E.A., Bruno-Blanch, L.E., Talevi, A.: Chapter 2 - Knowledge-Based Drug Repurposing: A Rational Approach Towards the Identification of Novel Medical Applications of Known Drugs. In: Ul-Haq, Z. and Madura, J.D. (eds.) Frontiers in Computational Chemistry. pp. 44–81. Bentham Science Publishers (2015)

29. Cheng, F., Desai, R.J., Handy, D.E., Wang, R., Schneeweiss, S., Barabási, A.-L., Loscalzo, J.: Network-based approach to prediction and population-based validation of in silico drug repurposing. Nat Commun. 9, 2691 (2018). 10.1038/s41467-018-05116-5

30. Mullins, J.G.L.: Drug repurposing in silico screening platforms. Biochemical Society Transactions. 50, 747–758 (2022). 10.1042/BST20200967

31. Hodos, R.A., Kidd, B.A., Shameer, K., Readhead, B.P., Dudley, J.T.: In silico methods for drug repurposing and pharmacology. WIREs Systems Biology and Medicine. 8, 186–210 (2016). 10.1002/wsbm.1337

32. Kumar, S., Kumar, S.: Chapter 6 - Molecular Docking: A Structure-Based Approach for Drug Repurposing. In: Roy, K. (ed.) In Silico Drug Design. pp. 161–189. Academic Press (2019)

33. Sohraby, F., Bagheri, M., Aryapour, H.: Performing an In Silico Repurposing of Existing Drugs by Combining Virtual Screening and Molecular Dynamics Simulation. In: Vanhaelen, Q. (ed.) Computational Methods for Drug Repurposing. pp. 23–43. Springer, New York, NY (2019)

34. McConachie, S.M., Wilhelm, S.M., Kale-Pradhan, P.B.: New direct-acting antivirals in hepatitis C therapy: a review of sofosbuvir, ledipasvir, daclatasvir, simeprevir, paritaprevir, ombitasvir and dasabuvir. Expert Review of Clinical Pharmacology. 9, 287–302 (2016). 10.1586/17512433.2016.1129272

35. Kumari, R., Kumar, R., Lynn, A.: g_mmpbsa—A GROMACS Tool for High-Throughput MM-PBSA Calculations. J. Chem. Inf. Model. 54, 1951–1962 (2014). 10.1021/ci500020m

36. Sun, M., Lv, S., Pan, Y., Song, Q., Ma, C., Yu, M., Gao, X., Guo, X., Wang, S., Gao, Z., Wang, S., Meng, Q., Zhang, L., Li, Y.: Discovery of Daclatasvir as a potential PD-L1 inhibitor from drug repurposing. Bioorganic Chemistry. 153, 107874 (2024). 10.1016/j.bioorg.2024.107874

37. Kim, S.: Exploring Chemical Information in PubChem. Current Protocols. 1, e217 (2021). 10.1002/cpz1.217

38. Knox, C., Wilson, M., Klinger, C.M., Franklin, M., Oler, E., Wilson, A., Pon, A., Cox, J., Chin, N.E. (Lucy), Strawbridge, S.A., Garcia-Patino, M., Kruger, R., Sivakumaran, A., Sanford, S., Doshi, R., Khetarpal, N., Fatokun, O., Doucet, D., Zubkowski, A., Rayat, D.Y., Jackson, H., Harford, K., Anjum, A., Zakir, M., Wang, F., Tian, S., Lee, B., Liigand, J., Peters, H., Wang, R.Q. (Rachel), Nguyen, T., So, D., Sharp, M., da Silva, R., Gabriel, C., Scantlebury, J., Jasinski, M., Ackerman, D., Jewison, T., Sajed, T., Gautam, V., Wishart, D.S.: DrugBank 6.0: the DrugBank Knowledgebase for 2024. Nucleic Acids Research. 52, D1265–D1275 (2024). 10.1093/nar/gkad976

39. Bogdanov, A., Bogdanov, A., Chubenko, V., Volkov, N., Moiseenko, F., Moiseyenko, V.: Tumor acidity: From hallmark of cancer to target of treatment. Front. Oncol. 12, (2022). 10.3389/fonc.2022.979154

40. Jo, S., Cheng, X., Lee, J., Kim, S., Park, S.-J., Patel, D.S., Beaven, A.H., Lee, K.I., Rui, H., Park, S., Lee, H.S., Roux, B., MacKerell Jr, A.D., Klauda, J.B., Qi, Y., Im, W.: CHARMM-GUI 10 years for biomolecular modeling and simulation. Journal of Computational Chemistry. 38, 1114–1124 (2017). 10.1002/jcc.24660

41. Huang, J., Rauscher, S., Nawrocki, G., Ran, T., Feig, M., de Groot, B.L., Grubmüller, H., MacKerell, A.D.: CHARMM36m: an improved force field for folded and intrinsically disordered proteins. Nat Methods. 14, 71–73 (2017). 10.1038/nmeth.4067

42. Abraham, M.J., Murtola, T., Schulz, R., Páll, S., Smith, J.C., Hess, B., Lindahl, E.: GROMACS: High performance molecular simulations through multi-level parallelism from laptops to supercomputers. SoftwareX. 1–2, 19–25 (2015). 10.1016/j.softx.2015.06.001

43. Genheden, S., Ryde, U.: The MM/PBSA and MM/GBSA methods to estimate ligand-binding affinities. Expert Opinion on Drug Discovery. 10, 449–461 (2015). 10.1517/17460441.2015.1032936

44. Saad, K.A., Eldawy, M.A., Elokely, K.M.: Studies of the symmetric binding mode of daclatasvir and analogs using a new homology model of HCV NS5A GT-4a. J Mol Model. 29, 25 (2022). 10.1007/s00894-022-05420-4

45. Langhans, B., Nischalke, H.D., Krämer, B., Hausen, A., Dold, L., van Heteren, P., Hüneburg, R., Nattermann, J., Strassburg, C.P., Spengler, U.: Increased peripheral CD4+ regulatory T cells persist after successful direct-acting antiviral treatment of chronic hepatitis C. Journal of Hepatology. 66, 888–896 (2017). 10.1016/j.jhep.2016.12.019

46. Guo, J., Yu, F., Zhang, K., Jiang, S., Zhang, X., Wang, T.: Beyond inhibition against the PD-1/PD-L1 pathway: development of PD-L1 inhibitors targeting internalization and degradation of PD-L1. RSC Medicinal Chemistry. 15, 1096–1108 (2024). 10.1039/D3MD00636K

47. Wang, T., Cai, S., Cheng, Y., Zhang, W., Wang, M., Sun, H., Guo, B., Li, Z., Xiao, Y., Jiang, S.: Discovery of Small-Molecule Inhibitors of the PD-1/PD-L1 Axis That Promote PD-L1 Internalization and Degradation. J. Med. Chem. 65, 3879–3893 (2022). 10.1021/acs.jmedchem.1c01682

48. Szereday, L., Meggyes, M., Berki, T., Miseta, A., Farkas, N., Gervain, J., Par, A., Par, G.: Direct-acting antiviral treatment downregulates immune checkpoint inhibitor expression in patients with chronic hepatitis C. Clin Exp Med. 20, 219–230 (2020). 10.1007/s10238-020-00618-3

49. Ganesan, A., Ahmed, M., Okoye, I., Arutyunova, E., Babu, D., Turnbull, W.L., Kundu, J.K., Shields, J., Agopsowicz, K.C., Xu, L., Tabana, Y., Srivastava, N., Zhang, G., Moon, T.C., Belovodskiy, A., Hena, M., Kandadai, A.S., Hosseini, S.N., Hitt, M., Walker, J., Smylie, M., West, F.G., Siraki, A.G., Lemieux, M.J., Elahi, S., Nieman, J.A., Tyrrell, D.L., Houghton, M., Barakat, K.: Comprehensive in vitro characterization of PD-L1 small molecule inhibitors. Sci Rep. 9, 12392 (2019). 10.1038/s41598-019-48826-6

50. Liu, M., Zhang, Y., Guo, Y., Gao, J., Huang, W., Dong, X.: A comparative study of the recent most potent small-molecule PD-L1 inhibitors: what can we learn? Med Chem Res. 30, 1230–1239 (2021). 10.1007/s00044-021-02728-3

51. Jafari, M., Guan, Y., Wedge, D.C., Ansari-Pour, N.: Re-evaluating experimental validation in the Big Data Era: a conceptual argument. Genome Biology. 22, 71 (2021). 10.1186/s13059-021-02292-4

52. Pillai, M., Wu, D.: Validation approaches for computational drug repurposing: a review. AMIA Annual Symposium Proceedings. 2023, 559 (2024)

53. Experimental validation, anyone? Nat Comput Sci. 3, 361–361 (2023). 10.1038/s43588-023-00462-x

