## Supplementary material of Daclatasvir, a symmetric drug for an anti-symmetric target for "Daclatasvir, a symmetric drug for an anti-symmetric target"

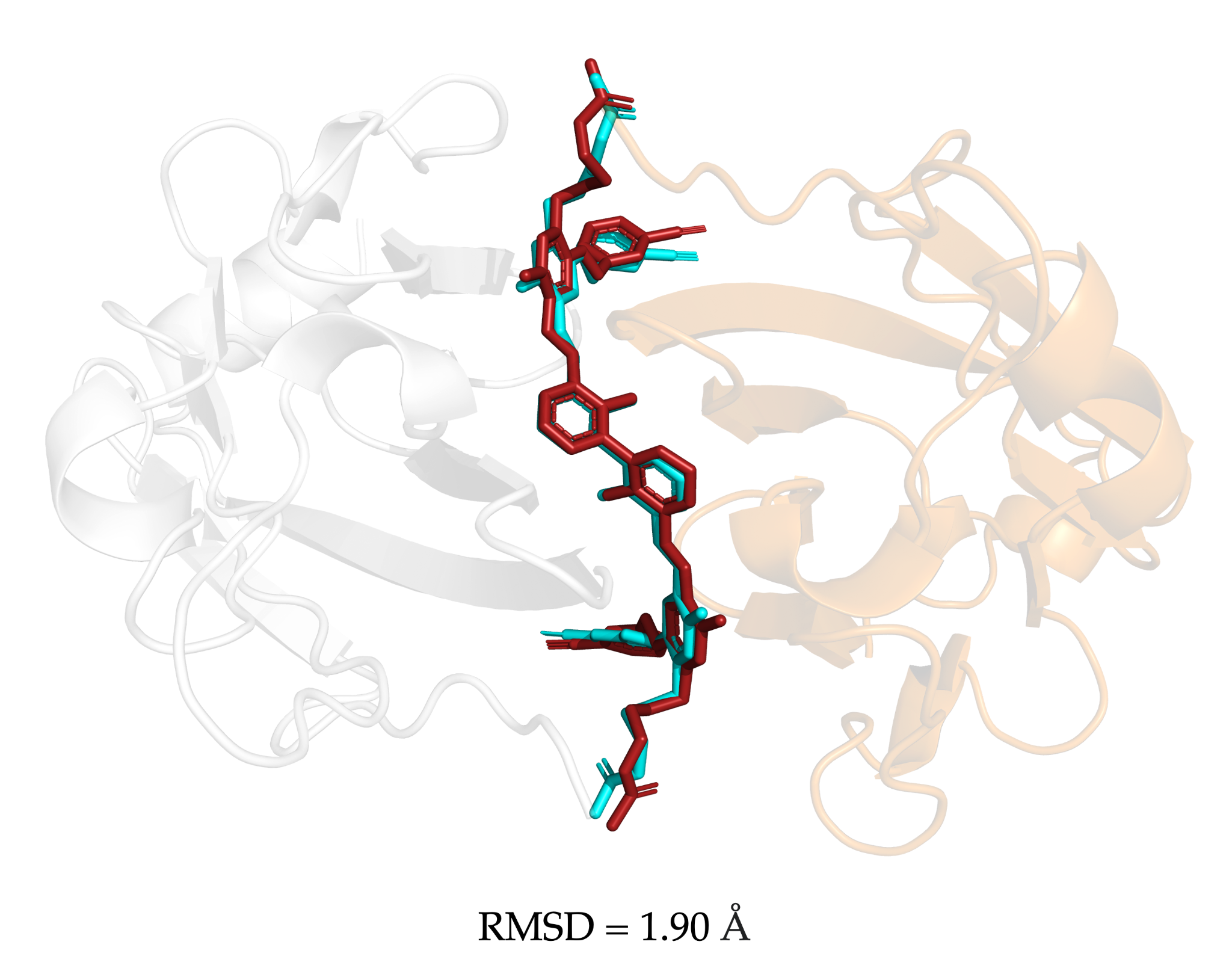

**Figure S1**. Superposition of co-crystallized (cyan) and redocked (red) ligands in the PD-L1 homodimer (PDB ID: 6RPG).

**Table S1**. Top-Ranked compounds from high-throughput molecular docking of FDA Database into the PD-L1 homodimer.

| **Ranking** | **Drug name** | **Score** | **Ranking** | **Drug name** | **Score** |
| --- | --- | --- | --- | --- | --- |
| **1** | Semaglutide | -266.74 | **11** | Triptorelin | -220.44 |
| **2** | Pramlintide | -264.74 | **12** | Lypressin | -220.15 |
| **3** | Daptomycin | -254.42 | **13** | Bleomycin | -220.01 |
| **4** | Tannic acid | -253.31 | **14** | Tetracosactide | -218.62 |
| **5** | Bivalirudin | -236.78 | **15** | Ledipasvir | -216.91 |
| **6** | Mivacurium | -232.59 | **16** | Etelcalcetide | -216.16 |
| **7** | Ubiquinol | -230.51 | ***17*** | *Daclatasvir* | *-215.96* |
| **8** | Cefoperazone | -228.76 | **18** | Gonadorelin | -211.43 |
| **9** | Fondaparinux | -225.84 | **19** | Isavuconazonium | -210.42 |
| **10** | Micafungin | -223.42 | **20** | Histrelin | -208.06 |

**Table S2.** Binding free energy* of PD-L1 homodimer complexes.

|  |  | **ΔH** | **-TΔS** | **ΔG** |
| --- | --- | --- | --- | --- |
| **Ligand A** | R1 | -64.03 ± 5.88 | 33.55 ± 14.02 | -30.48 ± 15.21 |
|  | R2 | -74.72 ± 6.16 | 31.30 ± 15.72 | -43.42 ± 16.88 |
|  | R3 | -60.89 ± 7.67 | 42.14 ± 22.15 | -18.75 ± 23.44 |
|  | *Average* | *-66.55 ± 3.82* | *35.66 ± 10.19* | *-30.88 ± 10.88* |
| **Ligand B** | R1 | -43.67 ± 5.35 | 32.13 ± 11.05 | -11.54 ± 12.28 |
|  | R2 | -41.07 ± 5.86 | 17.58 ± 6.00 | -23.50 ± 8.39 |
|  | R3 | -46.55 ± 7.92 | 33.55 ± 14.18 | -13.00 ± 16.24 |
|  | *Average* | *-43.76 ± 3.74* | *27.75 ± 6.32* | *-16.01 ± 7.34* |
| **Daclatasvir** | R1 | -34.97 ± 4.69 | 7.24 ± 2.96 | -27.73 ± 5.54 |
|  | R2 | -30.46 ± 5.16 | 5.69 ± 2.89 | -24.77 ± 5.91 |
|  | R3 | -34.15 ± 5.61 | 19.85 ± 10.22 | -14.29 ± 11.66 |
|  | *Average* | *-33.19 ± 2.98* | *10.93 ± 3.68* | *-22.26 ± 4.73* |

*All values are reported in kcal/mol.
